# Non-canonical roles of caspase-4 and caspase-5 in heme driven- IL-1β release and cell death

**DOI:** 10.1101/2020.02.28.969899

**Authors:** Beatriz E. Bolívar, Alexandra N. Brown, Brittany A. Rohrman, Chloé I. Charendoff, Vanda Yazdani, John D. Belcher, Gregory M. Vercellotti, Jonathan M. Flanagan, Lisa Bouchier-Hayes

## Abstract

Excessive release of heme from red blood cells is a key pathophysiological feature of several disease states including bacterial sepsis, malaria, and sickle cell disease. This hemolysis results in an increased level of free heme that has been implicated in the inflammatory activation of monocytes, macrophages, and endothelium. Here, we show that extracellular heme engages the human inflammatory caspases, caspase-1, caspase-4, and caspase-5, resulting in the release of IL-1β. Heme-induced IL-1β release was further increased in macrophages from patients with sickle cell disease. In human primary macrophages, heme activated caspase-1 in an inflammasome-dependent manner, but heme-induced activation of caspase-4 and caspase-5 was independent of canonical inflammasomes. Furthermore, we show that both caspase-4, and caspase-5 are essential for heme-induced IL-1β release, while caspase-4 is the primary contributor to heme-induced cell death. Together, we have identified that extracellular heme acts as a damage associated molecular pattern (DAMP) that can engage canonical and non-canonical inflammasome activation as a key mediator of inflammation in macrophages.

**Footnotes:** Funding for the project includes NIH/NIDDK T32DK060445 (BEB, BAR), NIH/NIDDK F32DK121479 (BEB), NIH/NIGMS R01GM121389 (LBH), NIH/NHLBI R01HL114567 (JDB, GMV), NIH/NHLBI R01-HL136415 (JMF), and CPRIT-RP180672, NIHCA125123 and NIHRR024574 (Cytometry and Cell Sorting Core at Baylor College of Medicine)

**Key Points:**

1. Heme induces oligomerization of caspase-1, caspase-4, and caspase-5.
2. Heme-induced IL-1β release requires both caspase-4 and caspase-5.
3. Caspase-4 alone contributes to heme-induced cell death.

## Introduction

The interactions between inflammatory caspases and inflammasomes are critical for preventing uncontrolled inflammation and for mediating appropriate inflammation under infectious and sterile conditions. Inflammasomes are multi-protein complexes that provide the platform for recruitment and activation of inflammatory caspases and are essential for cellular inflammatory responses (1). The inflammatory caspases include the human caspases-1, -4, and -5 and murine caspase-11 (1). This subset of the broader caspase protease family does not mediate apoptosis, but specifically regulates inflammation by facilitating the activation and release of the pro-inflammatory cytokines interleukin (IL)-1β and IL-18 (2). While the essential nature of inflammatory caspases in pathogen clearance is well established, their role in sterile inflammation (inflammation in the absence of infection) is less clear. Sterile inflammation occurs when non-pathogenic inflammatory stimuli activate inflammasomes. These stimuli are known as damage-associated molecular patterns (DAMPs) and are generally endogenous signals released by dying cells. This type of inflammation is important for wound healing and tissue regeneration but, if unchecked, can contribute to tissue damage associated with conditions like ischemic stroke, myocardial infarction, and neurodegeneration (3). Despite the importance of DAMPs for triggering inflammation, the endogenous signaling molecules that trigger sterile inflammation are not fully resolved (4).

Heme has the features of a DAMP because it is released following red blood cell destruction and triggers an inflammatory response. Extracellular hemoglobin and heme are highly pro-oxidant molecules that are assiduously scavenged by haptoglobin and hemopexin, respectively. Excessive hemolysis can saturate and deplete the haptoglobin and hemopexin systems, resulting in free heme with strong pro-inflammatory capabilities (5). Heme has been shown to activate caspase-1 in mouse macrophages via assembly of the NOD-like receptor family pyrin domain containing 3 (NLRP3) inflammasome (6). Heme has also been shown to activate toll-like receptor (TLR) 4 in murine endothelial cells to activate NFκB (7). For IL-1β release to proceed, two signals are needed. Signal 1 activates NFκB to induce pro-IL-1β expression and expression of additional inflammasome proteins including NLRP3, in a process known as priming. Signal 2 provides an intracellular signal that induces inflammasome assembly and caspase-1 activation that cleaves pro-IL-1β to its mature form, which is released from the cell (8). Heme is naturally taken up and recycled by macrophages, providing a physiological intracellular Signal 2 (44). Due to its ability to activate both caspase-1 and TLR4 and to be internalized by macrophages, extracellular heme has the properties of a DAMP that could potentially provide both Signal 1 and Signal 2 to initiate an effective inflammatory response.

Excessive release of heme from red blood cells is a key feature of several pathological states, including sepsis, malaria, and sickle cell disease (SCD). SCD is the most prevalent inherited blood disorder, affecting approximately 100,000 Americans and millions worldwide (9). The clinical manifestations of SCD arise from a complex pathophysiology including chronic hemolytic anemia, increased susceptibility to infection, and vaso-occlusive events (10). Chronically elevated heme levels induce the inflammatory activation of monocytes, macrophages, and the endothelium (7, 11, 12). This inflammation can result in vaso-occlusion, acute chest syndrome, and organ damage (5, 7, 13). Heme-induced activation of monocytes and macrophages contributes to these severe complications through release of inflammatory cytokines, like IL-1β, that trigger endothelial activation, upregulation of adhesion factors, and vaso-occlusion (14, 15). Indeed, in a study of children with SCD, patients having a vaso-occlusive pain crisis demonstrated elevated levels of proinflammatory cytokines IL-1β, IL-6, IL-10, tumor necrosis factor (TNF)-α, and free heme (16). The role of the inflammatory caspases in heme-induced inflammation in the context of hemolytic disorders such as SCD has not been well studied.

Mice deficient in caspase-1 or the inflammasome proteins NLRP3 or apoptosis-associated speck-like protein containing a CARD (ASC) survive following hemolysis induced by a lethal dose of phenylhydrazine (6). Thus, caspase-1 is essential for an effective inflammatory response to hemolysis. However, the roles of caspase-4 and caspase-5 in this process are unknown. Caspase-4 and caspase-5 are the human orthologues of murine caspase-11. Caspase-4, -5, and -11 have been shown to be activated by intracellular LPS independent of inflammasomes and they each have been shown to cleave the pore-forming protein gasdermin D (GSDMD) (17-19). Cleavage of GSDMD allows its N-terminal fragment to insert in the plasma membrane forming a pore predicted to be 180 Å in diameter (20). This pore is of sufficient size to allow release of mature IL-1β, but also permits influx of ions leading to cell swelling and a necrotic form of cell death called pyroptosis (21-23). Although caspase-1 can also cleave GSDMD, blocking caspase-1-dependent cleavage delays, but does not inhibit, pyroptosis (17). Therefore, current thinking is that caspase-1 cleaves pro-IL-1β and pro-IL-18, while caspase-4, caspase-5, and caspase-11 cleave GSDMD to induce pyroptosis, allowing active cytokine release. It has been proposed that cytokine release and pyroptosis cannot be uncoupled (24). However, contradicting this theory, some reports show living cells releasing IL-1β (25, 26). Here, we show that heme induces inflammasome-dependent caspase-1 activation and inflammasome-independent caspase-4 and caspase-5 activation. Furthermore, we show that both caspase-4 and caspase-5 are essential for IL-1β release and caspase-4 contributes to heme-induced cell death.

## Materials and Methods

### Chemicals and antibodies

The following antibodies were used: anti-Caspase-1 (D7F10 from Cell Signaling Technology, Danvers, MA, USA), anti-Caspase-4 (4450 from Cell Signaling Technology, Danvers, MA, USA), anti-Caspase-5 (D3G4W from Cell Signaling Technology, Danvers, MA, USA); anti-actin (C4 from MP Biomedicals); anti-GSDMD (G7422 from Sigma-Aldrich, St. Louis, MO, USA), anti-GSDMD N-term (E7H9G from Cell Signaling Technology, Danvers, MA, USA), anti-IL-1β (MAB601, R&D Systems, Inc., Minneapolis, MN, USA), anti-IL-1β cleaved, Asp116 (PA5-105048 from Thermo Fisher, Waltham, MA, USA). All cell culture media reagents were purchased from Thermo Fisher (Waltham, MA, USA). Ultrapure LPS (from E. coli O111:B4) was purchased from Invivogen (San Diego, CA, USA). Unless otherwise indicated, all other reagents were purchased from Sigma-Aldrich (St. Louis, MO, USA).

### Plasmids

The pBiFC.VC155 and pBiFC.VN173 plasmids encoding C1-Pro, C4-Pro, or C5-Pro were described previously (27). Single mutations were introduced using QuikChange Site-Directed Mutagenesis Kit (Agilent Technologies, Santa Clara, CA, USA). The bicistronic vector consists of C1-Pro VC and C1-Pro VN linked with a 2A peptide. Silent mutations were introduced into the second C1-Pro nucleotide sequence to prevent the sequence recombining out during cloning due to the presence of two identical C1 Pro sequences. The sequence was generated by IDT (Coralville, IA, USA) and cloned into pRRL-MND-MCS-2A-mCherry-2A-Puro. pDsRed-mito was purchased from Clontech (Takara, Mountain View, CA, USA). Each construct was verified by sequencing.

### Preparation and culture of primary human monocyte-derived macrophages

Whole blood samples were obtained from patients with SCD who attend our hematology clinic as part of their routine care. The study was approved by the Baylor College of Medicine Institutional Review Board, and informed consent was obtained from all participants, or legal guardian (if participants were minor). To isolate peripheral blood mononuclear cells, whole blood obtained from healthy blood donors or SCD patients was separated using the Ficoll Paque (GE Healthcare, Pittsburg, PA, USA) gradient protocol (28). CD14+ monocytes were isolated from peripheral blood mononuclear cells (PBMC) using magnetic bead selection (Miltenyi Biotec, San Diego, CA, USA). To differentiate cells into macrophages, monocytes were seeded at 5×10^6^ -1×10^7^/10 cm dish in RPMI-1640 medium supplemented with FCS (10% (v/v)), glutamax (2 mM), and Penicillin/ Streptomycin (50 I.U./50 µg/ml) and granulocyte-macrophage colony-stimulating factor (GM-CSF; 50 ng/mL). Cells were allowed to adhere overnight and culture media was exchanged the following day for fresh GM-CSF-supplemented media. Media was exchanged every 2-3 days and were considered fully differentiated at 7 days, as determined by morphology. To differentiate into M2 macrophages, media was exchanged and cells were incubated for additional 24 hours in media supplemented with IL-4 (50 ng/mL). For the matched M1 macrophages, cells were incubated in fresh GM-CSF supplemented media for an additional day.

### Heme preparation and administration

Heme solution was prepared immediately before use by solubilizing 3.3 mg of porcine hemin, (oxidized version of heme) in 100 µL of NaOH solution (0.1-1.0 M). The mixture was vortexed for 5 min in the dark. 900 µL of serum-free RPMI-1640 media was added to the resulting solution and vortexed for an additional 5 min in the dark followed by filtration through a sterile 0.22 µm spin filter. The term “heme” is used generically to refer to both heme and hemin. Unless otherwise indicated, cells were treated with heme in the presence of 0.1% FBS in the culture media to prevent components present in FBS sequestering and inhibiting heme. Heme is rapidly recycled by circulating macrophages, therefore, to mimic the transient exposure of cells to heme and to limit its toxicity, where indicated, cells were exposed to heme for 1 h followed by addition of an equal volume of culture media containing 10% FBS to inactivate the heme.

### Cell culture and generation of cell lines

THP-1 cells were grown in RPMI medium containing FBS (10% (v/v)), glutamax (2 mM), and Penicillin/ Streptomycin (50 I.U./50 µg/ml). MCF-7 cells were grown in Dulbecco’s Modified Essential Medium (DMEM) containing FBS (10% (v/v)), L-glutamine (2 mM), and Penicillin/ Streptomycin (50 I.U./50 µg/ml). *CASP1*-deficient THP-1 cells were purchased from Invivogen (San Diego, CA, USA). Caspase-4 and caspase-5 were deleted from THP-1 cells using an adaptation of the CRISPR/Cas9 protocol described in (29). Protospacer sequences for each target gene were identified using the CRISPRscan scoring algorithm (www.crisprscan.org (30)). DNA templates for sgRNAs were made by PCR using pX459 plasmid containing the sgRNA scaffold sequence and using the following primers: ΔCASP4(40) sequence: TTAATACGACTCACTATAGGGAAACAACCGCACACGCCgttttagagctagaaatagc; ΔCASP5(46) sequence: TTAATACGACTCACTATAGGTCCTGGAGAGACCGCACAgttttagagctagaaatagc; ΔCASP5(53) sequence: TTAATACGACTCACTATAGGTCAAGGTTGCTCGTTCTAgttttagagctagaaatagc; universal reverse primer: AGCACCGACTCGGTGCCACT. sgRNAs were generated by *in vitro* transcription using the Hiscribe T7 high yield RNA synthesis kit (New England Biolabs, Ipswich, MA, USA). Purified sgRNA (0.5 μg) was incubated with Cas9 protein (1 μg, PNA Bio, Newbury Park, CA, USA) for 10 min at room temperature. THP-1 cells were electroporated with the sgRNA/Cas9 complex using the Neon transfection system (Thermo Fisher Scientific, Waltham, MA, USA) at 1600V, 10ms, and 3 pulses. For the caspase-5 deficient cell line, two sgRNA were selected at either end of the gene to delete the intervening region. Deletion of *CASP4* or *CASP5* was confirmed by western blot or by PCR. Single cell clones were generated by single cell plating of the parental cell line. Gene deletion in the single cell clones was confirmed by sequencing.

### Transient Transfection and siRNA

For transfection of hMDM, 1 × 10^5^ cells were transfected with the appropriate plasmid combinations or plasmid/siRNA combinations using the Neon Transfection System (Thermo Fisher, Waltham, MA, USA) and a 10 µL Neon tip at 1000 V, 40 ms, and 2 pulses. Cells were transfected with amounts of the relevant expression plasmids as described in the figure legends. A total of 4 wells were transfected for every plasmid and incubated in 200 µL of antibiotic-free media. After 1 h, 200 µL of complete growth media containing Penicillin/ Streptomycin (50 I.U./50 µg/ml) was added. Expression was allowed for 24 h and media was exchanged for fresh media prior to treatment. Control siRNAs were siCyclophilin B (ON-TARGETplus SMARTpool from Dharmacon). ASC and NLRP3 siRNAs were ON-TARGETplus SMARTpool (Dharmacon Inc, Lafayette, CO, USA). 1 × 10^5^ MCF-7 cells were transfected with appropriate plasmid combinations using Lipofectamine 2000 transfection reagent (Thermo Fisher Scientific, Waltham, MA, USA) according to manufacturer’s instructions.

### Microscopy

Cells were imaged using a spinning disk confocal microscope (Zeiss, Thornwood, NY, USA), equipped with a CSU-X1A 5000 spinning disk unit (Yokogowa Electric Corporation, Japan), multi laser module with wavelengths of 458 nm, 488 nm, 514 nm, and 561 nm, and an Axio Observer Z1 motorized inverted microscope equipped with a precision motorized XY stage (Carl Zeiss MicroImaging, Thornwood, NY, USA). Images were acquired with a Zeiss Plan-Neofluar 40× 1.3 NA or 64× 1.4 NA objective on an Orca R2 CCD camera using Zen 2012 software (Zeiss, Thornwood, NY, USA). Cells were plated on dishes containing coverslips (Mattek Corp. Ashland, MA, USA) coated with poly-D-lysine hydrobromide 24 h prior to treatment. For time-lapse experiments, media on the cells was supplemented with Hepes (20 mM) and 2-mercaptoethanol (55 μM). Cells were allowed to equilibrate to 37°C in 5% CO_2_ prior to focusing on the cells.

### Image analysis

Images were analyzed by drawing regions around individual cells and then computing average intensity the pixels for each fluor using Zen 2012 software (Zeiss, Thornwood, NY, USA). Data were scaled by the following formula: scaled point = (Max − *x*)/MaxDifference, where Max equals the maximum value in the series, *x* equals the point of interest, and MaxDifference equals the maximum minus the minimum value in the series.

### ELISA IL-1β Measurements

THP-1 cells were plated at 1×10^6^ cells/ mL and differentiated into macrophages by 24 h incubation in the presence of phorbol 12-myristate 13-acetate (PMA; 10 ng/mL) followed by 24 hour incubation in RPMI medium. Cells were polarized into M1 macrophages by 20 h incubation with human interferon (IFN)-γ (PeproTech, 20 ng/mL) and ultrapure LPS (10 pg/mL) (31). Cells were washed prior to treatment as indicated in the Figure Legends. IL-1β concentration in harvested clarified supernatants was measured with the IL-1β Duoset ELISA Kit (R&D Systems, Minneapolis, MN, USA) according to the manufacturer’s instructions.

### Flow cytometry and measurement of cell death

Cells were treated as indicated and collected by centrifugation. Cells were washed with PBS and resuspended in 100 µl of PBS supplemented with 5% FCS, 0.5% BSA, 2 mM EDTA and 1 µl of 7-AAD (Thermo Fisher, Waltham, MA, USA). 7-AAD-positive cells were quantitated by flow cytometry and analyzed with FloJo software (FloJo LLC., Ashland, OR, USA).

### Immunoblotting

Cells were treated as indicated. Cells were lysed in IP-lysis buffer (50 mM Tris pH 7.4, 150 mM NaCl, 0.1% SDS, 1% NP-40 containing protease inhibitors (cOmplete Mini Protease Inhibitor Cocktail). Protein concentration was determined by BCA assay (Thermo Fisher, Waltham, MA, USA). 25-35 µg total protein (lysate) or 30-60 µg culture media (supernatant) was resolved by SDS-PAGE and transferred onto 0.45 µm nitrocellulose membrane (Thermo Fisher, Waltham, MA, USA), and immunodetected using appropriate primary and peroxidase-coupled secondary antibodies (GE Healthcare Life Sciences, Pittsburg, PA, USA). Proteins were visualized by West Pico and West Dura chemiluminescence Substrate (Thermo Fisher, Waltham, MA, USA).

### Statistical Analysis

Statistical comparisons were performed using two-tailed Student’s t test or 1-way ANOVA calculated using Prism 6.0 (Graph Pad) software.

## Results

### Heme-induced IL-1β release from human macrophages requires priming

It has been previously shown that heme can induce IL-1β release in LPS-primed murine bone marrow derived macrophages and that heme can activate TLR4 to induce NFκB activation in endothelial cells (6, 7). This suggests that heme can provide both the priming signal (Signal 1) required to induce pro-IL-1β expression and the inflammasome activating signal (Signal 2) required to induce inflammasome-dependent caspase-1 activation. To test this in human macrophages, we isolated CD14+ monocytes from peripheral blood from healthy donors and differentiated them into macrophages with granulocyte-macrophage colony-stimulating factor (GM-CSF). We exposed the macrophages to heme in the presence or absence of prior priming with LPS. Treatment with heme in non-primed macrophages induced a modest increase in IL-1β. However, in order for heme to induce maximal IL-1β release, prior priming with LPS was required. Treatment with LPS alone resulted in similar IL-1β release to heme alone (Figure 1A). These experiments were carried out in 0.1% FBS in the culture media. This is because components present in serum including hemopexin and albumin bind to heme with high affinity and prevent its uptake into cells (32-35). To show that the IL-1β release was specific to heme and nothing else present in the formulation, we increased the amount of serum in the culture media. As little as 1% FBS concentration in the cellular media was sufficient to block heme-induced IL-1β release (Figure 1B), indicating that the IL-1β release from macrophages that we detected was due to the specific effects of heme.

**Figure 1:**
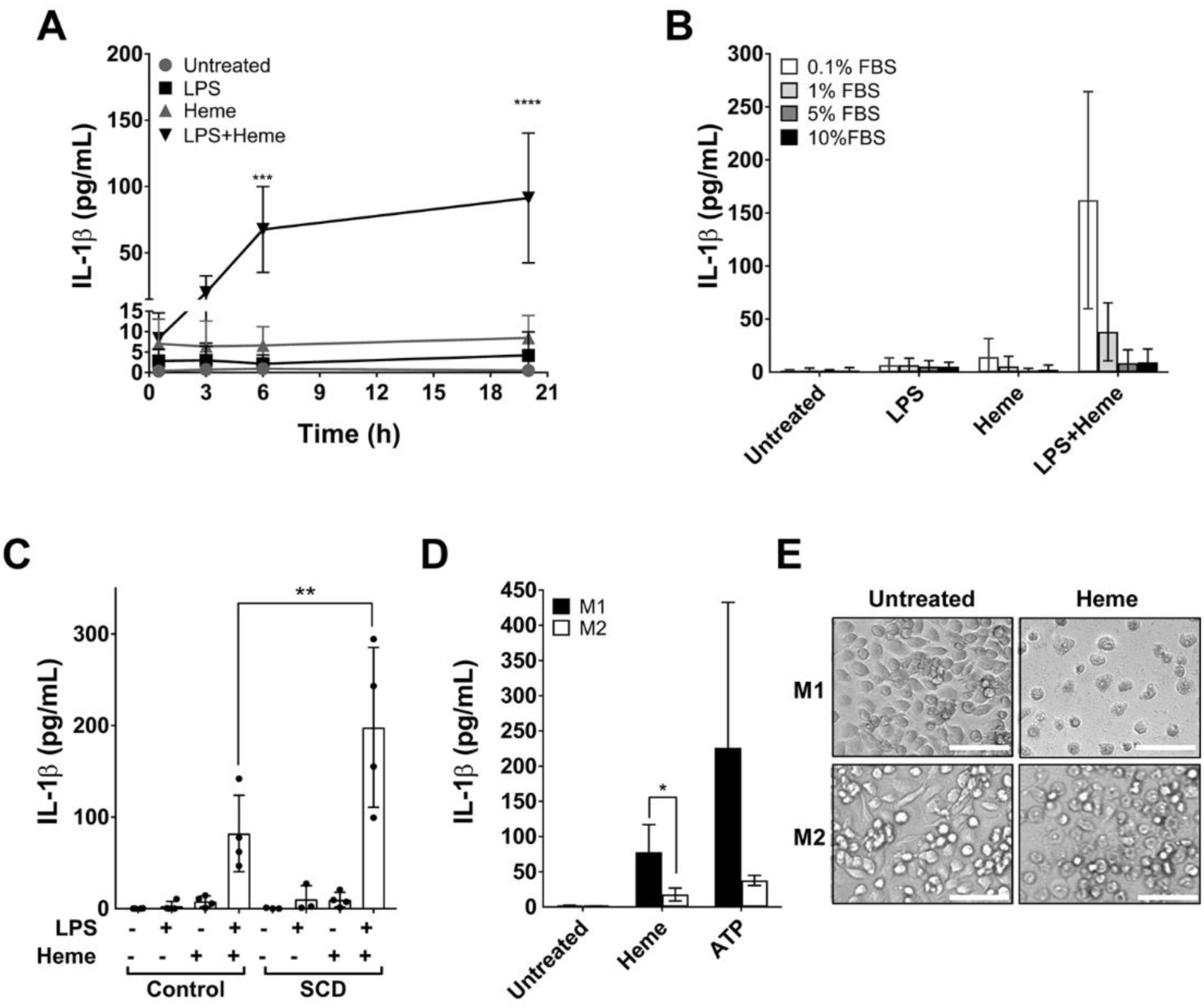
Heme induces IL-1β release that is increased in SCD patients. **(A)** CD14+ monocytes were isolated from 5 healthy donors and differentiated into macrophages using GM-CSF for 7 days. When fully mature, cells were primed with or without LPS (100 ng/mL) for 3 h, washed and treated with or without heme (50 µM) in 0.1% FBS. Mature IL-1β levels were measured in cellular supernatants by ELISA at the indicated times. Error bars represent standard deviation of 5 independent experiments. ***p<0.001, ****p<0.0001 calculated by 1-way ANOVA. **(B)** GM-CSF-differentiated human macrophages from healthy donors were primed with or without LPS for 3 h (100 ng/mL), followed by treatment with or without heme (50 µM) in 0.1, 1, 5, or 10% FBS. After 20 h, IL-1β concentration was measured in cultured supernatants by ELISA. Error bars represent standard deviation of 4 independent biological replicates. **(C)** GM-CSF-differentiated human macrophages were isolated from healthy donors (control) or patients with sickle cell disease (SCD) and primed with or without LPS (100 ng/mL) for 3 h followed by treatment with or without heme (50 µM) in 0.1% FBS. IL-1β concentration was measured in cultured supernatants at 20 h by ELISA. Error bars represent standard deviation of 4 control and 4 SCD samples across 4 independent experiments. **p<0.01 calculated by 1-way ANOVA. **(D)** CD14+ monocytes were isolated from 3-5 healthy donors and differentiated in GM-CSF for 7 days. On day 7, macrophages were polarized either into the M2-like phenotype with IL-4 (50 ng/mL) or into the M1-like phenotype with GM-CSF for an additional 24 h. On day 8, both groups were washed and primed with LPS (100 ng/mL) for 3 h followed by treatment with or without heme (50 µM) in 0.1% FBS or ATP (5 mM) in 10% FBS. IL-1β concentration was measured in cultured supernatants at 20 h by ELISA. Error bars represent standard deviation of 3-5 independent experiments. *p<0.05 calculated by Student’s t test. **(E)** Representative phase-contrast images of M1 and M2 macrophages from (D) are shown. Scale bar represents 100 µm.

Sickle cell macrophages are constantly exposed to high levels of extracellular heme due to persistent hemolysis of sickled red blood cells (11). To determine the effect of this on IL-1β release, we compared human monocyte-derived macrophages (hMDM) from healthy donors to hMDM isolated from patients with SCD. We noted that heme-treated hMDM from patients with SCD were more sensitive to heme-induced IL-1β release relative to the control hMDM (Figure 1C). However, priming with LPS was required for strong IL-1β release in both groups. Together, these results suggest that, while heme is able to activate caspase-1 to induce IL-1β release, it is not sufficient to prime the cells for IL-1β release.

It has been shown that macrophages from sickle mice express higher levels of M1 markers (36). To investigate IL-1β release by heme in the two extreme macrophage subtypes, we polarized the CD14+ monocytes towards M1 (pro-inflammatory) by treating with GM-CSF for seven days or towards M2 (anti-inflammatory) by treating with IL-4 for a further day after the incubation with GM-CSF. LPS-primed M1 or M2 hMDM were stimulated with heme or with ATP. We showed lower levels of IL-1β release in M2 compared to M1 hMDM in response to both heme and ATP (Figure 1D). In response to heme, M1 macrophages underwent cell death with a necrotic morphology. In contrast, M2 macrophages remained viable with an intact plasma membrane but they appeared more flattened and granular than the unstimulated cells (Figure 1E). Together, these results suggest that SCD macrophages have increased inflammatory caspase activity, resulting in increased IL-1β activation and that this may be due, in part, to the increased proportion of M1 macrophage population in patients with SCD.

### Caspase-1, caspase-4, and caspase-5 are activated by heme in the absence of priming

Given that heme induces IL-1β release in macrophages, we next investigated the ability of heme to activate caspase-1 and the other human inflammatory caspases, caspase-4 and caspase-5. Caspase-1 is activated upon proximity-induced dimerization following recruitment to inflammasomes (1, 37). The recruitment of caspase-1 to an inflammasome is mediated by interactions between specific protein interaction domains in the inflammasome proteins. The caspase recruitment domain (CARD) in ASC or in NLR family CARD containing 4 (NLRC4) binds to the CARD in the prodomain of caspase-1 to recruit it to the inflammasome and facilitate induced proximity of the caspase (38, 39). To measure the induced proximity of each caspase, we used caspase bimolecular fluorescence complementation (BiFC) (40). BiFC uses non-fluorescent fragments of the yellow fluorescent protein, Venus (“split Venus”) that can associate to reform the fluorescent Venus complex when fused to interacting proteins (41). When the CARD-containing prodomain of caspase-1 is fused to each half of split Venus, recruitment of caspase-1 to inflammasomes, and the subsequent induced proximity results in enforced association of the two Venus halves (27). Thus, Venus fluorescence (BiFC) acts as a read-out for caspase induced proximity, the proximal step for activation. We transiently expressed the prodomain of caspase-1 (C1-Pro, aa 1-102) fused to each of the split Venus fragments, Venus C (aa 155-239) and Venus N (aa 1-173) in hMDM (Figure 2A). Following exposure to heme, we noted a significant increase in the proportion of cells that became Venus positive. Surprisingly, and unlike the requirement for IL-1β release, this induction of caspase-1 BiFC did not require prior priming with LPS. When we transfected macrophages with the caspase-4 prodomain (C4-Pro) or caspase-5 prodomain (C5-Pro) BiFC pairs, we observed a similar pattern, where caspase-4 and caspase-5 BiFC were induced by heme, independent of LPS priming (Figure 2B,C). This suggests that caspase-1, caspase-4, and caspase-5 are each activated by heme in macrophages. When we analyzed the appearance and localization of the fluorescent complexes produced by heme in each case, we noticed some differences (Figure 2D). The caspase-1 complex appeared as a single green punctum typical of ASC specks and of ASC-induced caspase-1 BiFC that we previously reported (27, 42). Caspase-4 and caspase-5 BiFC appeared as a series of punctate spots located throughout the cytoplasm of the cell that did not co-localize with mitochondria. This indicates that there may be different mechanisms of action for caspase-1 compared to caspase-4 or caspase-5.

**Figure 2:**
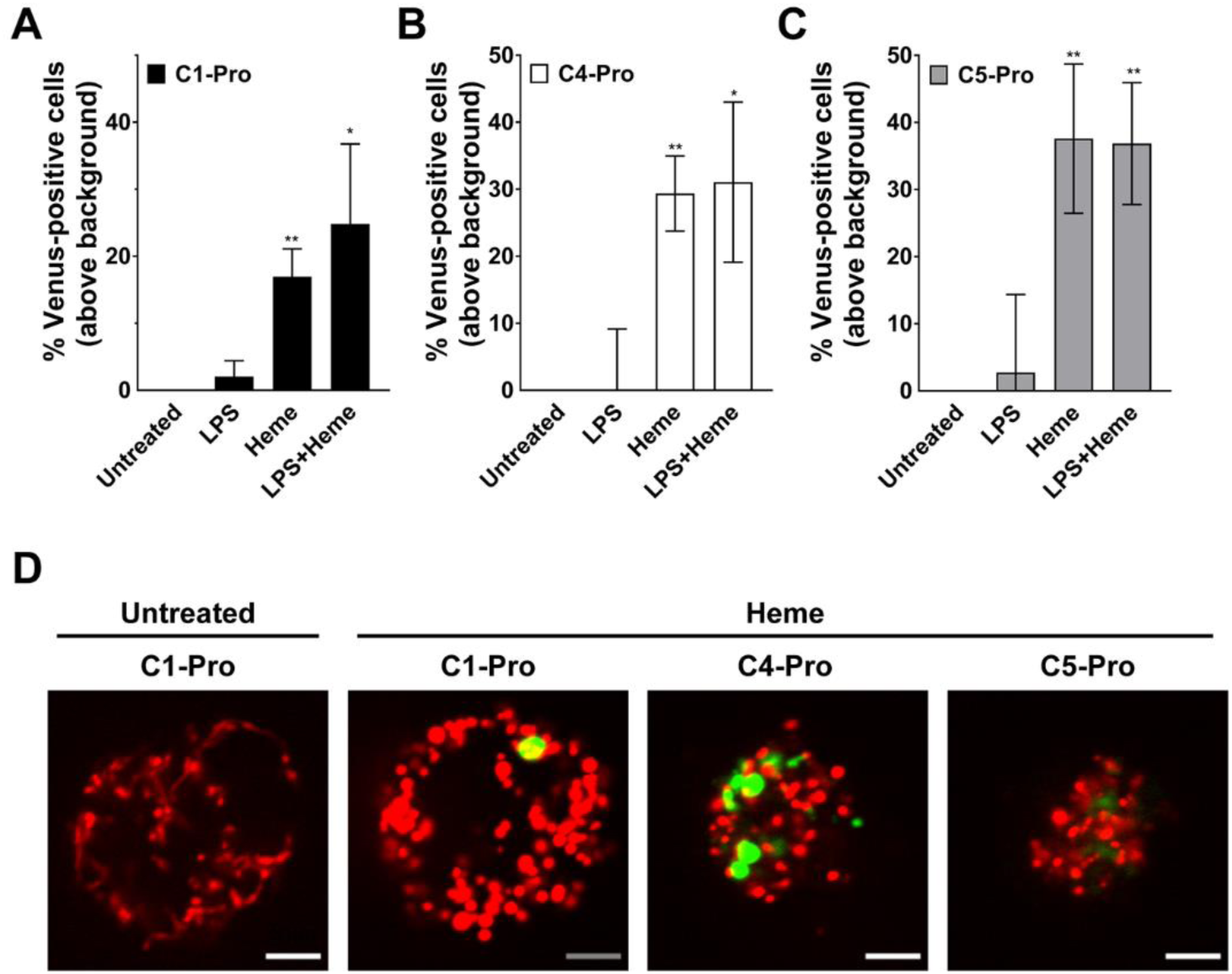
Heme activates the inflammatory caspases. GM-CSF-differentiated human macrophages isolated from healthy donors were transfected with the C1-Pro VC (300 ng) and C1-Pro VN (300 ng) **(A)**; C4-Pro VC (500 ng) and of C4-Pro VN (500 ng) **(B)**; or C5-Pro VC (1000 ng) and C5-Pro VN (1000 ng) **(C)**; along with dsRedmito (50 ng) as a reporter for transfection. 24 h after transfection, cells were treated with or without LPS (100 ng/mL) for 3 h followed by treatment with or without heme (50 µM) in 0.1% FBS. After 1 h, FBS was reconstituted to 5% to inhibit extracellular heme. Cells were assessed for the percentage of dsRed-positive transfected cells that were Venus-positive at 20 h, determined from a minimum of 300 cells per well. Results are represented as percent Venus-positive cells over background (untreated cells). Error bars represent standard deviation of four independent experiments. *p< 0.05; **p< 0.01; calculated by Student’s t test. **(D)** Representative images show caspase BiFC in green and mitochondria in red. Scale bar represents 10 µm.

### Caspase-5 activation is impaired in M2 macrophages

Next, we used the BiFC system to measure inflammatory caspase activation in M1 and M2 macrophages. We polarized macrophages into M1 and M2 as before and transfected donor-matched M1 and M2 cells with each of the inflammatory caspase BiFC pairs. Following exposure to heme, caspase-1, caspase-4, and caspase-5 BiFC was increased as before in M1 hMDM (Figure 3A). Interestingly, while caspase-1 and caspase-4 BiFC was induced to the same extent in M1 and M2 macrophages, the M2 macrophages had impaired heme-induced caspase-5 BiFC. To explore this further, we measured caspase-1, caspase-4, and caspase-5 expression in M1 and M2 macrophages by immunoblot (Figure 3B). Endogenous caspase-1 and caspase-4 were similarly expressed in both subtypes and their expression was unchanged with the addition of heme. Caspase-5 expression was readily detected in M1 macrophages, but was lowly expressed in unstimulated M2 macrophages. Heme did not induce caspase-5 expression in the M2 subgroup, but pre-treatment with LPS restored the level of caspase-5 to the level detected in M1 cells. We hypothesized that the low level of endogenous caspase-5 in M2 hMDM was responsible for the inability of these cells to induce caspase-5 BiFC. To test this, we compared caspase-5 BiFC induced by heme in M1 and M2 hMDM with and without LPS priming. Priming of M2 macrophages with LPS restored their ability to induce the caspase-5 BiFC complex (Figure 3C). This suggests that full length endogenous caspase-5 is required to form the caspase-5 activation complex in response to heme.

**Figure 3:**
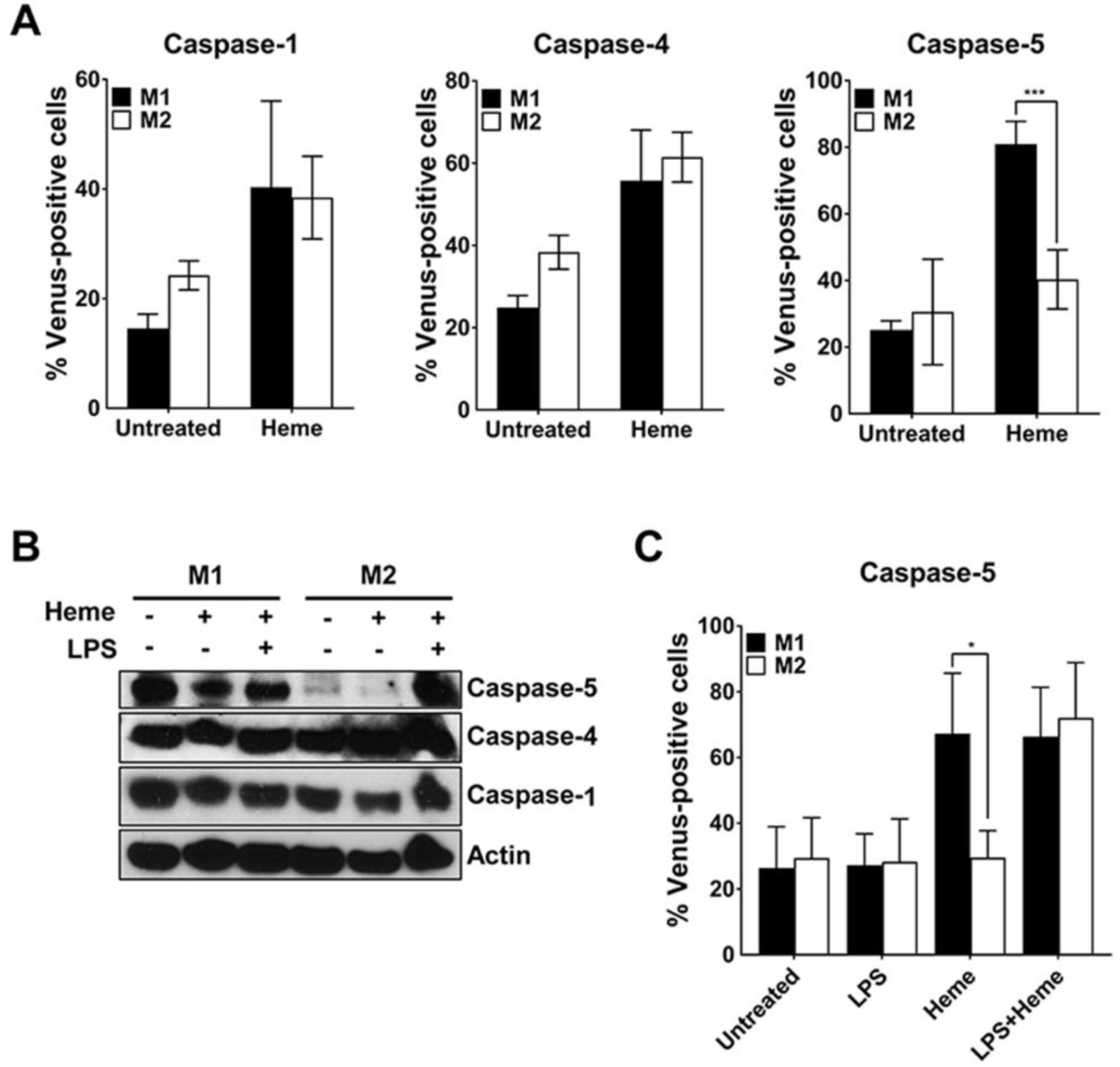
Heme-induced caspase-5 activation and expression is reduced in M2 macrophages. **(A)** GM-CSF-differentiated human macrophages isolated from healthy donors were polarized into M1 or M2 macrophages and transfected with the C1-Pro BiFC pair (300 ng of each), the C4-Pro BiFC pair (500 ng of each), or the C5-Pro BiFC pair (1000 ng of each) along with dsRedmito (50 ng) as a reporter for transfection. 24 h after transfection, cells were treated with or without heme (50 µM) in 0.1% FBS. After 1 h, FBS was reconstituted to 5% to inhibit extracellular heme. Cells were assessed for the percentage of dsRed-positive transfected cells that were Venus-positive at 20 h, determined from a minimum of 300 cells per well. Error bars represent standard deviation of three independent experiments. ***p=0.001; calculated by Student’s t test. **(B)** Cells were polarized to M1 or M2 macrophages as in (A) and treated with or without LPS (100 ng/mL) for 3 h followed by heme (50 µM) in 0.1% FBS. After 1 h, FBS was reconstituted to 5% to inhibit extracellular heme and 20 h later cell lysates were immunoblotted for caspase-1, caspase-4, caspase-5, or actin as a loading control. **(C)** Cells were polarized to M1 or M2 macrophages and transfected with the C5-Pro BiFC pair as in (A). Transfected hMDMs were treated with or without LPS (100 ng/mL) for 3 h followed by heme (50 µM) in 0.1% FBS. After 1 h, FBS was reconstituted to 5% to inhibit extracellular heme and 20 h later cells were assessed for the percentage of dsRed-positive transfected cells that were Venus-positive, determined from a minimum of 300 cells per well. Error bars represent standard deviation of three independent experiments. *p<0.05; calculated by Student’s t test.

### Heme activates caspase-4 and caspase-5 independently of canonical inflammasome interactions

Intracellular LPS has been shown to bind and induce oligomerization of caspase-4, caspase-5, or caspase-11 independently of inflammasomes (19). The oligomerization is mediated by CARD clustering, which leads to induced proximity and caspase dimerization. Since heme can promote caspase-4 and caspase-5 induced proximity and is naturally taken up and recycled by macrophages, we reasoned that heme may represent an intracellular trigger for non-canonical inflammatory caspase activation (activation independent of known inflammasomes). To test this, we investigated if heme-induced caspase activation requires interactions with inflammasomes. Recruitment of caspases to inflammasomes is dependent on an intact CARD. The aspartate residue (D59) in caspase-1 is essential for the ASC-caspase-1 interaction and mutation of this D59 residue blocks ASC-induced caspase-1 BiFC (27, 43). We previously showed that ASC overexpression does not induce caspase-4 or caspase-5 BiFC but NLRC4 does (27). We modified the conserved CARD-binding residue found in caspase-4 (D59) and caspase-5 (D117), and showed that disruption of this residue also blocked caspase-4 and caspase-5 BiFC triggered by overexpressed NLRC4 (Supplemental Figure S1A, B). Using the CARD-disrupting mutant in caspase-1 (D59R), caspase-4 (D59R), and caspase-5 (D117R), we tested if heme-induced inflammatory caspase BiFC is independent of canonical inflammasome interactions. We transfected hMDM with the caspase-1, caspase-4 or caspase-5 Pro BiFC pairs or with the corresponding CARD-disrupting mutant BiFC pairs. Following exposure to heme, caspase-1 (D59R) BiFC was significantly decreased compared to the wild type caspase-1 reporter (Figure 4A). This indicates that heme triggers recruitment of caspase-1 to the inflammasome for its subsequent activation. In contrast, heme-induced caspase-4 and capase-5 BiFC was not changed when the CARD mutant BiFC reporters were expressed (Figure 4A). This suggests that heme activates caspase-4 and caspase-5 independently of canonical inflammasome interactions.

**Figure 4:**
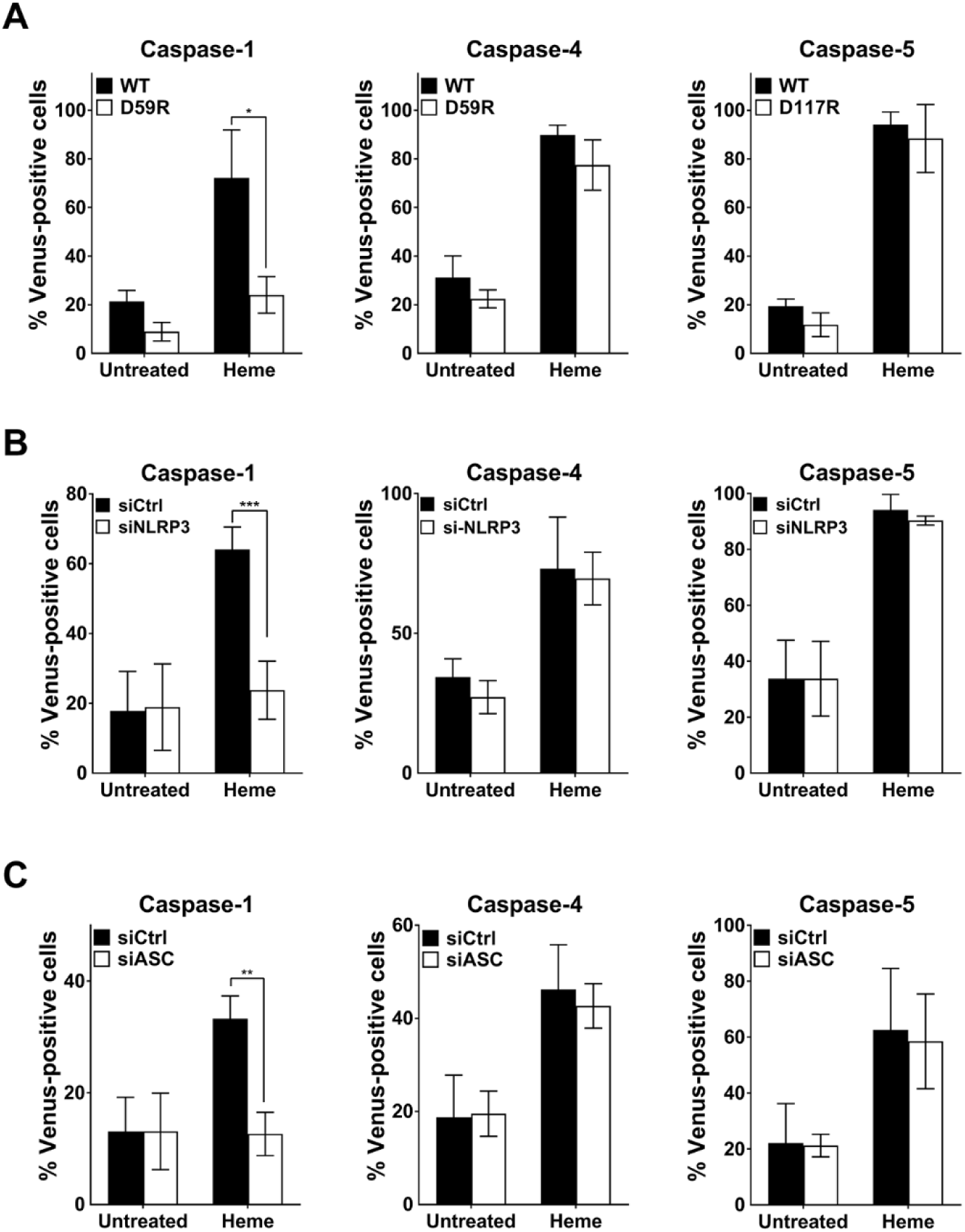
Heme activates caspase-4 and caspase-5 independently of canonical inflammasome interactions. **(A)** GM-CSF-differentiated human macrophages isolated from healthy donors were transfected with the C1-Pro BiFC pair (300 ng of each), the D59R mutant C1-Pro BiFC pair (300 ng of each), the C4-Pro BiFC pair (500 ng of each), the D59R mutant C4-Pro BiFC pair (500 ng of each), the C5-Pro BiFC pair (1000 ng of each), or the D117R mutant C5-Pro BiFC pair (1000 ng of each) along with dsRedmito (50 ng) as a reporter for transfection. 24 h after transfection, cells were treated with or without heme (50 µM) in 0.1% FBS. After 1 h, FBS was reconstituted to 5% to inhibit extracellular heme. Cells were assessed for the percentage of dsRed-positive transfected cells that were Venus-positive at 20 h, determined from a minimum of 300 cells per well. Error bars represent standard deviation of four independent experiments. *p<0.05; calculated by Student’s t test. **(B)** GM-CSF-differentiated human macrophages isolated from healthy donors were transfected with the C1-Pro BiFC pair (300 ng of each), the C4-Pro BiFC pair (500 ng of each), or the C5-Pro BiFC pair (1000 ng of each), along with dsRedmito (50 ng) as a reporter for transfection with either siRNA against NLRP3 or a control siRNA (7.5 pmol). 24 h after transfection, cells were treated with or without heme (50 µM) in 0.1% FBS. After 1 h, FBS was reconstituted to 5% to inhibit extracellular heme. Cells were assessed for the percentage of dsRed-positive transfected cells that were Venus-positive at 20 h, determined from a minimum of 300 cells per well. Error bars represent standard deviation of three independent experiments. ***p<0.001; calculated by Student’s t test. **(C)** GM-CSF-differentiated human macrophages isolated from healthy donors were transfected with the C1-Pro BiFC pair (300 ng of each), the C4-Pro BiFC pair (500 ng of each), or the C5-Pro BiFC pair (1000 ng of each), along with dsRedmito (50 ng) as a reporter for transfection with either siRNA against ASC or a control siRNA (7.5 pmol). 24 h after transfection, cells were treated with or without heme (50 µM) in 0.1% FBS. After 1 h, FBS was reconstituted to 5% to inhibit extracellular heme. Cells were assessed for the percentage of dsRed-positive transfected cells that were Venus-positive at 20 h, determined from a minimum of 300 cells per well. Error bars represent standard deviation of three independent experiments. **p<0.01; calculated by Student’s t test.

To confirm these results, we used siRNA to silence NLRP3, the inflammasome receptor that has been shown to be required for heme-induced caspase-1 activation (6). We similarly silenced the inflammasome adaptor protein ASC. ASC is essential to the assembly of most characterized inflammasomes including the NLRP1, NLRP3, and AIM2 inflammasomes (1). It has been shown to be dispensable for the NLRC4 inflammasome, but ASC enhances NLRC4 inflammasome activity (44, 45). Therefore, if heme induces inflammatory caspase activation via the NLRP3 or alternative inflammasome complex, loss of ASC is predicted to block this activity. Si-RNA-mediated silencing of both NLRP3 and ASC in hMDM reduced heme-induced caspase-1 BiFC to background levels (Figure 4B, C Supplemental Figure S1C). In contrast, NLRP3 or ASC depletion had no effect on the levels of caspase-4 or caspase-5 BiFC induced by heme (Figure 4B, C). Together, these results show that caspase-4 and caspase-5 are activated by heme independently of canonical inflammasomes.

### Heme-induced cytokine release is dependent on caspase-4 and caspase-5

Having identified caspase-4 and caspase-5 induced proximity following heme exposure, we set out to explore the functional outcomes of this mechanism. We used CRISPR/Cas9 to delete caspase-4 and caspase-5 from the monocytic THP-1 cell line. We compared caspase-4 and caspase-5-deficient cells to parental THP-1 cells and to caspase-1 deficient THP-1 cells (purchased from Invivogen). To most closely represent the M1 phenotype of primary macrophages, cells were first treated with PMA followed by incubation with IFNγ and a low concentration of LPS (10 pg/mL) to polarize them to an M1-like state (31). The cells were then primed for 3 h with a higher concentration of LPS (100 pg/mL) followed by exposure to heme for a 1 h pulse. IL-1β release was induced by heme in LPS-primed M1 THP-1 cells to a similar level as we observed in M1 hMDM (∼80 pg/mL). As expected, caspase-1 deficiency completely blocked the release. Deficiency of either caspase-4 or caspase-5 potently blocked heme-induced IL-1β release. These results show that both caspase-4 and caspase-5 are required for IL-1β release induced by heme and that one caspase does not functionally replace for the action of the other.

Caspase-1, caspase-4, and caspase-5 have been shown to cleave GSDMD (18), the proposed mechanism by which inflammatory caspases induce both pyroptosis and release of mature cytokines. We therefore investigated GSDMD cleavage in M1 polarized THP-1 cells. In cells primed with LPS followed by exposure to heme, we did not detect levels of cleaved GSDMD over background in the cell lysates but we did detect it in the cellular supernatants (Figure 5B, lane 5). This is consistent with reports that N terminal of GSDMD (GSDMD-N) is released from cells when it is cleaved (21, 22). The amount of GSDMD-N that was produced by heme in LPS-primed cells was considerably lower than that of the positive control nigericin, a bacterial potassium ionophore and potent inducer of the NLRP3 inflammasome (Figure 5B, lane 6) (46, 47). Caspase-1 deficiency did not reduce the amount of GSDMD-N produced by heme treatment. In cells deficient in caspase-4, the amount of GSDMD-N detected in the cellular supernatant was lower compared to wild type cells but there was more detected in the lysate of heme treated LPS-primed cells. In cells deficient in caspase-5, a slight reduction in GSDMD-N was evident in supernatant. Therefore, the cleavage of GSDMD was not completely blocked in the absence of either caspase. Cleavage of GSDMD was not noticeably impaired in any of the caspase-deficient cells treated with nigericin. Together these results suggest that there is redundancy between caspase-4 and caspase-5 with respect to GSDMD cleavage. We also probed for IL-1β in the lysates and supernatants of each cell line. We were not able to detect the p17 IL-1β fragment in either fraction, but we detected a larger intermediate fragment running at around 25 kD in the supernatants of LPS-primed M1 THP-1 cells treated with heme. Production of this fragment was blocked in caspase-4 deficient and caspase-5 deficient cells and substantially reduced in the caspase-1 deficient cells. This is consistent with the reduction of IL-1β release detected by ELISA in these cells in Figure 5A. Importantly, the fact that we can still detect GSDMD-N in the heme-treated caspase-4 and caspase-5 deficient cells but IL-1β release is blocked in both of these cell types suggests that caspase-4 and caspase-5 have roles in regulating caspase-1 that are independent of GSDMD cleavage.

**Figure 5:**
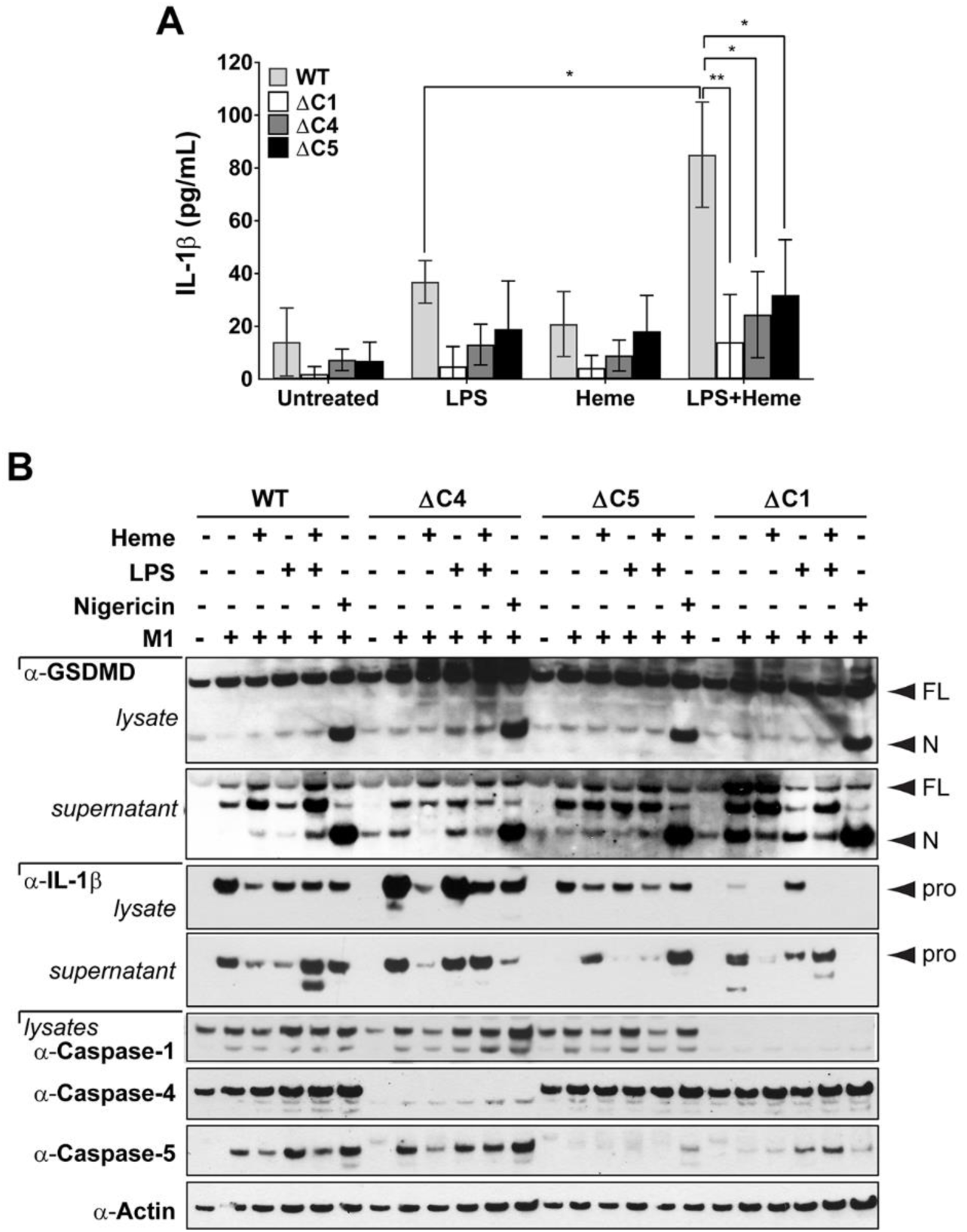
Heme-induced IL-1β release requires caspase-4 and caspase-5. **(A)** THP-1 cells (WT), THP-1 cells deficient in caspase-1 (ΔC1, generated by shRNA), caspase-4 (ΔC4, generated by CRISPR/Cas9), or caspase-5 (ΔC5, generated by CRISPR/Cas9) were treated with PMA (10 ng/mL) for one day and allowed to recover for an additional day followed by incubation with IFNγ (20 ng/mL) and LPS (10 pg/mL) for 20 h to polarize them into an M1 phenotype. Cells were treated with or without LPS (100 pg/mL) for 3 h followed by heme (50 µM) in 0.1% FBS. After 1 h, FBS was reconstituted to 5% to inhibit extracellular heme. IL-1β concentration was measured in cultured supernatants by ELISA at 20 h. Error bars represent standard deviation of 4 independent experiments. *p<0.05; **p<0.01; calculated by Student’s t test. **(B)** Unstimulated or M1 polarized THP-1 cells (M1) were treated with or without LPS (100 pg/mL) for 3 h followed by treatment with or without heme (50 µM) for 1 h in 0.1% FBS. 20 h later, cell lysates and culture supernatants were immunoblotted for the indicated proteins or actin as a loading control. FL, full length; N, GSDMD-N-term; pro, pro-IL-1β. Results are representative of three independent experiments.

### Heme-induced cell death is mediated in part by caspase-4

When we probed the cells in Figure 5B for pro-IL-1β, we noted a higher amount of pro-IL-1β in the lysates of caspase-4 deficient cells compared to wild type, caspase-5 deficient, or caspase-1 deficient cells. We reasoned that this may be due to the cells being protected from cell death in the absence of caspase-4. In addition, in the caspase-1 deficient cells treated with heme, all of the pro-IL-1β was detected in the supernatant and not the lysate suggesting cell lysis. This suggested that loss of caspase-1 has minimal effects on heme-induced cell death. To investigate the requirement of each caspase for heme-induced cell death, we first determined the level of death that was caspase-dependent. We treated LPS-primed or unprimed THP-1 cells with heme in the presence or absence of the pan-caspase inhibitor qVD-OPH (Figure 6A). Heme treatment of both primed and unprimed THP-1 cells induced substantial cell death as measured by 7-AAD uptake. Caspase inhibition significantly blocked this death but, notably, the reduction in cell death was not complete and approximately 30% of the cells were killed by heme in a caspase-independent manner. To determine the specific contribution of each inflammatory caspase to this death, we measured heme-induced cell death in caspase-1, caspase-4, and caspase-5-deficient THP-1 cells compared to wild type THP-1 cells. We measured cell death at two time points: an early time point at 6 h and a later time point at 20 h. At 6 h, heme did not induce a significant amount of death above background and there was no significant difference in the amount of death induced in the caspase-deficient cell lines (Figure 6B). At the 20 h time point, loss of caspase-4 significantly decreased heme-induced cell death, while loss of caspase-1 or caspase-5 had no effect (Figure 6C). This suggests that, among the inflammatory caspases, heme-induced caspase-4 activation is the primary effector of heme-induced cell death. Similar to the effect of total caspase inhibition, caspase-4 loss did not completely block the death and approximately 50% of the cells still died. This confirms that caspase-independent cell death mechanisms also contribute to heme-induced cell death.

**Figure 6.**
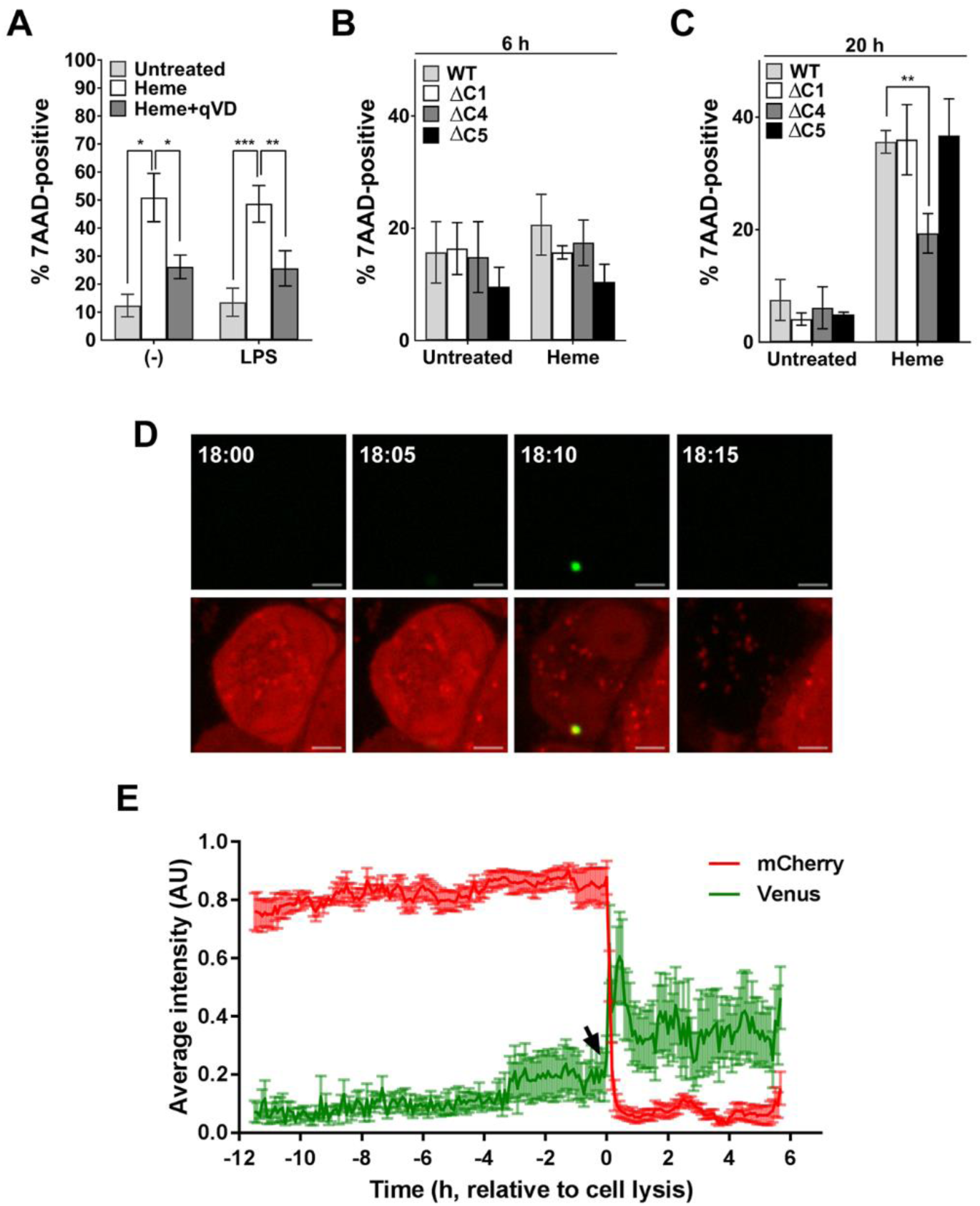
Caspase-4 contributes to heme-induced cell death. **(A)** THP-1 cells were treated with or without LPS (100 ng/mL) for 3 h in the presence or absence of qVD-OPH (5 µ) followed by treatment with or without heme (50 µM) for 1 h in 0.1% FBS. After 1 h, FBS was reconstituted to 5% to inhibit extracellular heme. Cell death was assessed by flow cytometry for 7-AAD uptake 20 h later. Error bars represent standard deviation of 3 independent experiments. *p<0.05; **p<0.01; ***p<0.001 calculated by Student’s t test. **(B-C)** THP-1 cells or THP-1 cells deficient in the indicated caspases were treated with or without heme (50 µM) as in (A). Cell death was assessed by flow cytometry for 7-AAD uptake at 6 h (B) or 20 h (C). Error bars represent standard deviation of 3-4 independent experiments. **p<0.01; calculated by Student’s t test. **(D)** PMA-primed THP-1 cells stably expressing the C1-Pro BiFC pair were treated with heme in the presence of qVD-OPH (5 μM) to prevent cells from lifting off due to apoptosis. Images were taken by confocal microscopy every 5 min for 24 h. Frames from the time-lapse show representative cells undergoing BiFC (green) prior to cell lysis as measured by the loss mCherry (red). Scale bars represent 5 µm. **(E)** Graph of the cells from (D) that became Venus-positive is shown. Each point on the mCherry graph (*red)* is scaled and aligned to each point on the caspase-1 BiFC graph (*green*) that represents the average intensity of mCherry or Venus in the cell at 5 min intervals where time=0 is the point of onset of mCherry loss, representing cell lysis. Arrow shows the point of onset of caspase-1 BiFC immediately prior to cell lysis. Error bars represent SEM of 9 individual cells.

To investigate the kinetics of heme-induced caspase-1 activation relative to cell death, we used time-lapse confocal microscopy. For this, we generated THP-1 cells stably expressing the caspase-1 pro BiFC components. We designed a bicistronic construct, where the caspase-1 Pro-VC and caspase-1 Pro-VN are expressed in a single vector separated by the viral 2A self-cleaving peptide, similar to a caspase-2 reporter we previously described (48). This design ensures that the caspase-1 BiFC components are expressed at equal levels because they are translated from a single mRNA transcript. These cells also express a linked mCherry gene as a reporter for expression of the BiFC components. In addition, loss of mCherry fluorescence can be used to detect cell lysis. The caspase-1 BiFC complex was detected approximately 18 h following the addition of heme and its appearance was followed by lysis of the cell that was detected by loss of the mCherry protein upon cell rupture (Figure 6A, Supplemental Movie S1). Cell death occurred within 5 minutes of detection of the C1-Pro BiFC complex (Figure 6B). Thus, recruitment of caspase-1 to inflammasomes in response to heme immediately precedes cell death. It is important to note that this death occurred in the presence of the pan-caspase inhibitor qVD-OPH. The inclusion of qVD-OPH in the time-lapse experiment was to prevent apoptotic cell death that causes the cells to lift off the coverslip and impairs imaging as the cells move out of the focal plane. This, together with the results from Figure 6A suggests that there is a substantial caspase-independent component to heme-induced cell death. However, the close timing of caspase-1 recruitment to inflammasomes and cell lysis indicates that these events are mechanistically linked, which could imply that caspases contribute to the death in a manner that does not require their catalytic activity.

## Discussion

Our data show that heme activates caspase-1, caspase-4, and caspase-5. However, there are significant differences in the upstream requirements for activation, the localization of the activation complexes and the outcomes of activation of each caspase. We found that both caspase-4 and caspase-5, are required for heme-induced IL-1β release, while caspase-4 is the primary contributor to heme-induced cell death. Our results indicate that caspase-4 and caspase-5 have non-overlapping functions in heme-induced inflammation and caspase-1 activation. Together, these data underscore the important functions of inflammatory caspases in heme-induced sterile inflammation.

While SCD is primarily a disease characterized by anemia, many of its clinical complications are exacerbated by chronic inflammation. Sickle shaped red blood cells are more prone to hemolysis releasing excess heme into the blood stream that overwhelms the body’s heme scavenging systems. Consistent with this, we noted that heme-treated macrophages derived from patients with SCD released more IL-1β when compared to healthy controls. This suggests that SCD macrophages are more sensitive to heme-induced inflammation. Using peripheral blood mononuclear cell transcriptome profiles of patients with SCD compared to healthy controls, van Beers *et al* showed that the SCD cohort had higher expression of many markers of innate immunity including *TLR4, NLRP3, NLRC4, CASP1, IL-1* and *IL-18* (49). They also showed a positive correlation between *TLR4* expression and *IL-6* expression in SCD cells. Lanaro *et al* showed increased expression of *TNFα* and *IL-8* in SCD mononuclear cells (50). Because heme can activate TLR4 and, in turn, the transcription factor NFκB that controls transcription of many of these cytokines and proteins, it is likely that a lifetime exposure to heme in patients with SCD resulting from higher rates of hemolysis contributes to these elevated levels. We originally suspected that the increased IL-1β release we detected from SCD macrophages after heme exposure would be due to this lifetime exposure to heme, which would prime the cells for inflammasome activation by inducing expression of pro-IL-1β and other inflammasome components. However, heme alone was insufficient to trigger IL-1β release in macrophages from patients with SCD or from healthy donors indicating that an extra priming step is still required. Monocytes from patients with SCD have been shown to have increased levels of the macrophage markers CD14 and CD11b (14). In addition, liver macrophages from HbS sickle mice had increased surface expression of the M1 macrophage markers CD86, MHCII, iNOS, and IL-6 (11). This suggests that sickle macrophages are likely to skew to the M1-like pro-inflammatory phenotype. Treatment of HbS sickle mice with the heme scavenger hemopexin reverted the M1 polarization, indicating that heme is the inducer of M1 polarization (11). Thus, we propose that the increased proportion of activated and pro-inflammatory M1 macrophages in patients with SCD is the reason why the cells, once primed by LPS, are more sensitive to heme-induced IL-1β release rather than the effect of heme priming. Interestingly, heme alone was sufficient to induce caspase-1, caspase-4, and caspase-5 activation. Caspase-1 activation by heme has been reported to be dependent on the NLRP3 inflammasome (6) and we confirmed that observation here. Unlike the other inflammasome receptors, NLRP3 requires a priming step for activation (51, 52). Our results may suggest that heme can prime for NLRP3 expression, but does not provide enough of a priming signal to induce sufficient pro-IL-1β expression. Induction of NLRP3 expression has been shown to be dependent on reactive oxygen species (ROS) (8). Heme induces ROS and blocking ROS has been shown to inhibit both caspase-1 cleavage and IL-1β processing (6). Therefore, heme-induced ROS generation may be a mechanism by which heme can prime for NLRP3 expression but not for pro-IL-1β expression.

Our observation that heme-induced caspase-4 and caspase-5 induced proximity does not require LPS priming could be explained in one of two ways. Similar to caspase-1, heme may be sufficient to prime the cells for caspase activation. However, we also show that, in contrast to caspase-1, caspase-4 and caspase-5 induced proximity is activated by heme independently of CARD-inflammasome interactions and of the common inflammasome proteins NLRP3 and ASC. The lack of a priming requirement for caspase-4 and caspase-5 induced proximity is consistent with caspase activation that does not require an additional upstream receptor. Caspase-5 is lowly expressed in many cell types and its expression is induced by LPS and interferon (53). Despite this, primary M1-polarized macrophages express abundant caspase-5. M2 macrophages have reduced caspase-5 expression and display reduced caspase-5 induced proximity in response to heme. This observation may suggest that full length, endogenous caspase-5 is required to promote assembly of the caspase-5 signaling platform. Indeed, when we reconstituted caspase-5 expression in M2 macrophages, we were able to restore the levels of caspase-5 induced proximity. These results are consistent with studies that have shown that caspase-4 and caspase-5 are direct intracellular receptors for LPS (19). LPS binds to the CARD in these caspases to trigger oligomerization of caspase-4 or caspase-5 in the absence of any known inflammasome protein. Our data indicate that heme acts in an analogous fashion to induce caspase-4 or caspase-5 oligomerization either directly or through an as yet unidentified cytosolic mediator. Contrary to the report that LPS binding to caspase-4 or caspase-5 is CARD-mediated (19), our results suggest that full length caspase-5 is required for its scaffolding function because the BiFC construct contains only the CARD-containing prodomain, which is insufficient in the absence of endogenous caspase-5. Further supporting a mechanistic difference between heme-induced caspase-1 activation and heme-induced caspase-4 or caspase-5 activation, we observed differences in the size and localization of the caspase-4 and caspase-5 signaling complexes induced by heme compared to the single ASC-like speck observed for caspase-1. Heme is naturally taken up and recycled by macrophages (44), representing a physiological stimulus and, importantly, a trigger of sterile inflammation that directly engages caspase-4 and caspase-5. Thus, heme acts as a canonical stimulus by engaging inflammasome-dependent caspase-1 activation, and also as a non-canonical stimulus by activating caspase-4 and caspase-5 independent of canonical inflammasomes.

Caspase-4 and caspase-5 are often assumed to be redundant proteins that directly phenocopy caspase-11 in mice. Studies using a transgenic mouse generated to express human caspase-4 indicated that caspase-4 does not completely phenocopy caspase-11 and a recent study showed a broader reactivity of caspase-4 to LPS compared to caspase-11 (54, 55). In addition, humans are more sensitive to endotoxemia than rodents (40). These studies highlight the important variations between the human and murine innate immune response and suggest that the presence of an additional inflammatory caspase in humans may contribute to these differences. We show that caspase-4 and caspase-5 are both required for heme-induced IL-1β release, providing evidence of a co-operative regulation between these two caspases and caspase-1 rather than a redundant function, where one can replace for the other. In contrast, loss of caspase-4 or caspase-5 on their own only marginally reduced GSDMD cleavage. This suggests some redundancy of these caspases for GSDMD cleavage. Therefore, the ability of caspase-4 or caspase-5 to mediate IL-1β release cannot be explained by inhibition of GSDMD cleavage by either caspase. Caspase-11 has been proposed to regulate caspase-1 (56), but the exact mechanism of this is unclear. Further work is needed to determine how caspase-4 and caspase-5 cooperate to regulate caspase-1 activation in response to heme and other stimuli.

Despite the similar effects on GSDMD cleavage, caspase-5 did not impact cell death induced by heme in the same manner as caspase-4. This suggests that while caspase-4 and caspase-5 both contribute to GSDMD cleavage, caspase-4 is the main effector of cell death. Indeed, studies have shown IL-1β release from live macrophages indicating that the pore formed by GSDMD and pyroptosis are separable events (25, 26). The caspase-4-dependent death we observed that is induced by heme may even be independent of GSDMD, suggesting caspase-4-specific substrates that promote cell death. However, there is still a considerable amount of caspase-independent death that is induced by heme. These results suggest that additional death mechanisms are engaged by heme. Heme has been shown to induce RIPK3-dependent necrosis (57) and, therefore, this may be a contributing mechanism to the death we observed. RIPK3-dependent necrosis has been shown to limit pathogenic inflammation and associated tissue damage without impairing IL-1β release (58). Given that cell death occurs immediately following detection of the caspase-1 activation platform, it is possible that a similar mechanism occurs here, where the cell lysis is a result of concurrent activation of the RIPK3 pathway rather than caspase-dependent effects. Since K+ efflux is known to be necessary for NLRP3-induced caspase-1 activation (47), another explanation could be that the close timing of caspase-1 induced proximity and cell lysis indicates that caspase-1 is oligomerizing in response to early potassium efflux through a caspase-4/5-driven pore. The exact interplay between caspase-1 activation and caspase-4/-5 activation that is triggered by exposure to heme, and how this leads to cell death is something that requires further study.

The role of caspase-4 in heme-induced cell death may suggest that caspase-4 is a key contributor to tissue damage in humans. The association between caspase-11, excess pyroptosis, and inflammation-associated tissue damage is well established (56, 59). In a mouse model of SCD, the consequences of heme-induced inflammation is tissue damage that manifests as vaso-occlusion and death (7). Our results suggest that caspase-4, and not caspase-1 or caspase-5, is the primary contributor to heme-induced pyroptosis. Our model predicts that blocking caspase-4 would protect from tissue damage, while blocking caspase-1, caspase-4, or caspase-5 would protect from uncontrolled inflammation (fever, pain, etc.). Further exploration of the distinct roles of these caspases is required to fully understand how chronic inflammation can be controlled in hemolytic disorders.

## Acknowledgements

We thank the members of LBH’s lab past and present for helpful discussions and careful reading of the manuscript. We thank Doug Green and the members of his lab for valuable suggestions. This project was supported by the Cytometry and Cell Sorting Core at Baylor College of Medicine with the assistance of Joel M. Sederstrom.

## Abbreviations used in this article

ASC: apoptosis-associated speck-like protein containing a CARD
BiFC: bimolecular fluorescence complementation
CARD: caspase recruitment domain
DAMP: damage-associated molecular pattern
GM-CSF: granulocyte-macrophage colony-stimulating factor
GSDMD: gasdermin D
hMDM: human monocyte-derived macrophages
IFN: interferon
IL: interleukin
LPS: lipopolysaccharide
NLR: NOD-like receptor
NLRC: NLR family CARD containing
NLRP: NLR family pyrin domain containing
PBMC: peripheral blood mononuclear cells
SCD: sickle cell disease
TLR: toll like receptor

