## Supplemental material for "Non-canonical roles of caspase-4 and caspase-5 in heme driven- IL-1β release and cell death"

### *Supplemental Figures*

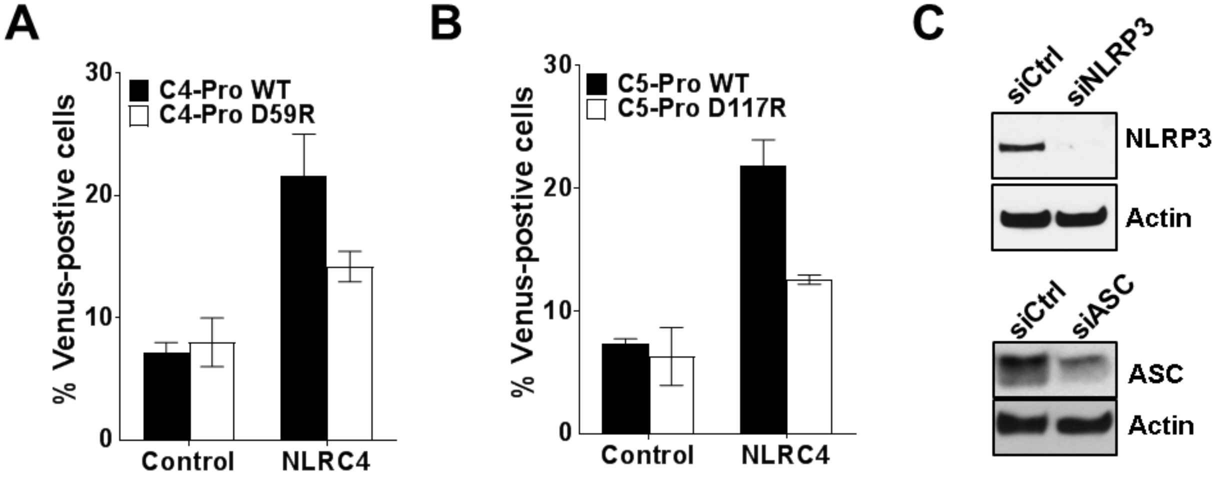


**Supplemental Figure S1: CARD mutants disrupt inflammasome induced inflammatory caspase BiFC.**

**(A)** MCF-7 cells were transiently transfected with C4-Pro VC (50 ng) and C4-Pro VN (50 ng) or the D59R mutant C4-Pro Venus pair (50 ng each) with or without an expression plasmid encoding NLRC4 (250 ng). All wells received dsRed-mito (10 ng) as a reporter for transfection. Cells were treated with qVD-OPH (5 μM) to prevent cell death. 48 h after transfection the percentage dsRed-mito-positive cells that were Venus-positive was determined from a minimum of 300 cells per well. Error bars represent standard deviation of two independent experiments. **(B)** MCF-7 cells were transiently transfected with C5-Pro VC (100 ng) and C5-Pro VN (100 ng) or the D177R mutant C5-Pro Venus pair (100 ng each) with or without an expression plasmid encoding NLRC4 (250 ng). All wells received dsRed-mito (10 ng) as a reporter for transfection. Cells were treated with qVD-OPH (5 μM) to prevent cell death. 48 h after transfection the percentage dsRed-mito-positive cells that were Venus-positive was determined from a minimum of 300 cells per well. Error bars represent standard deviation of two independent experiments. **(C)** GM-CSF-differentiated human macrophages isolated from healthy donors were transfected with either siRNA against NLRP3 (*upper*), ASC (*lower*) or a control siRNA (7.5 pmol). 48 h later cell lysates were immunoblotted for NLRP3, ASC or actin as a loading control. Results are representative of two independent experiments.

***Supplemental Movies***

**Supplemental Movie S1. Heme-induced inflammasome assembly immediately precedes cell death.** Movie of a representative PMA-primed THP-1 cell stably expressing the C1-Pro BiFC pair treated with heme in the presence of qVD-OPH (5 μM). Images were taken by confocal microscopy every 5 min for 24 h. The movie shows the cell undergoing BiFC (green) prior to cell lysis as measured by the loss mCherry (red). Scale bar represents 5 μm.
